# Heritability enrichment of specifically expressed genes identifies disease-relevant tissues and cell types

**DOI:** 10.1101/103069

**Authors:** Hilary K. Finucane, Yakir A. Reshef, Verneri Anttila, Kamil Slowikowski, Alexander Gusev, Andrea Byrnes, Steven Gazal, Po-Ru Loh, Caleb Lareau, Noam Shoresh, Giulio Genovese, Arpiar Saunders, Evan Macosko, Samuela Pollack, The Brainstorm Consortium, John R.B. Perry, Jason D. Buenrostro, Bradley E. Bernstein, Soumya Raychaudhuri, Steven McCarroll, Benjamin M. Neale, Alkes L. Price

## Abstract

Genetics can provide a systematic approach to discovering the tissues and cell types relevant for a complex disease or trait. Identifying these tissues and cell types is critical for following up on non-coding allelic function, developing ex-vivo models, and identifying therapeutic targets. Here, we analyze gene expression data from several sources, including the GTEx and PsychENCODE consortia, together with genome-wide association study (GWAS) summary statistics for 48 diseases and traits with an average sample size of 169,331, to identify disease-relevant tissues and cell types. We develop and apply an approach that uses stratified LD score regression to test whether disease heritability is enriched in regions surrounding genes with the highest specific expression in a given tissue. We detect tissue-specific enrichments at FDR < 5% for 34 diseases and traits across a broad range of tissues that recapitulate known biology. In our analysis of traits with observed central nervous system enrichment, we detect an enrichment of neurons over other brain cell types for several brain-related traits, enrichment of inhibitory over excitatory neurons for bipolar disorder but excitatory over inhibitory neurons for schizophrenia and body mass index, and enrichments in the cortex for schizophrenia and in the striatum for migraine. In our analysis of traits with observed immunological enrichment, we identify enrichments of T cells for asthma and eczema, B cells for primary biliary cirrhosis, and myeloid cells for Alzheimer's disease, which we validated with independent chromatin data. Our results demonstrate that our polygenic approach is a powerful way to leverage gene expression data for interpreting GWAS signal.

## INTRODUCTION

There are many diseases whose causal tissues or cell types are uncertain or unknown. Identifying these tissues and cell types is critical for developing systems to explore gene regulatory mechanisms that contribute to disease. In recent years, researchers have been gaining an increasingly clear picture of which parts of the genome are active in a range of tissues and cell types: for example, which parts of the genome are accessible, which enhancers are active, and which genes are expressed^1-3^. Combining this type of information with GWAS data offers the potential to identify causal tissues and cell types for disease.

Many different types of data characterizing tissue-and cell-type-specific activity have been analyzed together with GWAS data to identify disease-relevant tissues and cell types: histone marks^4-8^, DNase I hypersensitivity (DHS)^9-12^, eQTLs^3,13^, and gene expression data^14-17^ Of these data types, gene expression data (without genotypes or eQTLs) has the advantage of being available in the widest range of tissues and cell types. Therefore, methods for integrating gene expression data with GWAS data have the potential not only to identify system-level differences among traits—e.g., brain enrichment vs. immune enrichment—but also to obtain high resolution within a system—e.g., differentiating among brain regions or among immune cell types.

Indeed, previous work has shown that gene expression can be a useful source of information for identifying disease-relevant tissues and cell types from GWAS data. An initial application of the SNPsea algorithm^14,15^ analyzed a data set with gene expression in 249 immune cell types from mouse, together with genome-wide significant SNPs from GWAS of several immunological diseases, and reported disease-specific patterns of enrichment^13^. The DEPICT software^16^ includes a method for joint analysis of GWAS summary statistics with a large gene expression data set^18^, and has been used to identify enriched tissues for height^19^ and BMI^20^. In a recent study of migraine^17^, an analysis of genome-wide significant loci with expression data from the GTEx project identified cardiovascular and digestive/smooth muscle enrichments. These studies show that gene expression data are informative for disease-relevant tissues and cell types, and have led to biological insights about the diseases and traits studied. However, the methods applied in these studies restrict their analyses to subsets of SNPs that pass a significance threshold. To our knowledge, no previous study has modeled genome-wide polygenic signals to identify disease-relevant tissues and cell types from GWAS and gene expression data.

Here, we apply stratified LD score regression^7^, a method for partitioning heritability from GWAS summary statistics, to sets of specifically expressed genes to identify disease-relevant tissues and cell types across 48 diseases and traits with an average GWAS sample size of 169,331. We first analyze two gene expression data sets^3,16,18^ containing a wide range of tissues to infer system-level enrichments, recapitulating known biology. We also analyze chromatin data from the Roadmap Epigenomics and ENCODE projects^1,2^ across the same set of diseases and traits, and find that tissue-specific chromatin data can be used to validate results from tissue-specific gene expression. We then analyze gene expression data sets that allow us to achieve higher resolution within a system^3,21-23^, identifying enriched brain regions, brain cell types, and immune cell types for several brain-and immune-related diseases and traits; we validate several of our immune enrichments using independent chromatin data with a coarser representation of immune cells (13 cell types instead of 252). Our results underscore that a heritability-based framework applied to gene expression data allows us to achieve high-resolution enrichments, even for very polygenic traits.

## RESULTS

### Overview of methods

We analyzed the five gene expression data sets listed in Table 1, mapping mouse genes to orthologous human genes when necessary. To assess the enrichment of a focal tissue for a given trait, we follow the procedure described in Figure 1. We begin with a matrix of normalized gene expression values across genes, with samples from multiple tissues including the focal tissue. For each gene, we compute a t-statistic for specific expression in the focal tissue (Online Methods). We rank all genes by their t-statistic, and define the 10% of genes with the highest t-statistic to be the gene set corresponding to the focal tissue; we call this the set of specifically expressed genes, but we note that this includes not only genes that are strictly specifically expressed (i.e. only expressed in the focal tissue), but also genes that are weakly specifically expressed (i.e. higher average expression in the focal tissue). For a few of the data sets analyzed, we modified our approach to constructing the set of specifically expressed genes to better take advantage of the data available (Online Methods). We add 100kb windows on either side of the transcribed region of each gene in the set of specifically expressed genes to construct a genome annotation corresponding to the focal tissue. (The choice of the parameters 10% and 100kb is discussed in Online Methods; our results are robust to these choices (see below).) Finally, we apply stratified LD score regression^7^ to GWAS summary statistics to evaluate the contribution of the focal genome annotation to trait heritability (Online Methods). We jointly model the annotation corresponding to the focal tissue, a genome annotation corresponding to all genes, and the 52 annotations in the “baseline model”^7^ (including genic regions, enhancer regions, and conserved regions; see **Table S1**). A positive regression coefficient for the focal annotation in this regression represents a positive contribution of this annotation to trait heritability, conditional on the other annotations. We report regression coefficients, normalized by mean per-SNP heritability, together with a P-value to test whether the regression coefficient is significantly positive. Stratified LD score regression requires GWAS summary statistics for the trait of interest, together with an LD reference panel (e.g. 1000 Genomes^24^), and has been shown to produce robust results with properly controlled type I error^7^. We have released open source software implementing our approach, and have also released all genome annotations derived from the publicly available gene expression data that we analyzed (see URLs). We call our approach LD score regression applied to specifically expressed genes (LDSC-SEG).

**Table 1.** List of gene expression data sets used in this study. We analyzed five gene expression data sets: two (GTEx and Franke lab) containing a wide range of tissues and three (Cahoy, PsychENCODE, ImmGen) with more detailed information about a particular tissue.

| **Name** | **Organism** | **Tissues/cell types** | **Technology** |
| --- | --- | --- | --- |
| GTEx<sup>3</sup> | Human | 53 tissues/cell types | RNA-seq |
| Franke lab<sup>16,18</sup> | Human/mouse/rat | 152 tissues/cell types | Array |
| Cahoy<sup>21</sup> | Mouse | 3 brain cell types | Array |
| PsychENCODE<sup>22</sup> | Human | 2 neuronal cell types | RNA-seq |
| ImmGen<sup>23</sup> | Mouse | 292 immune cell types | Array |

**Figure 1:**
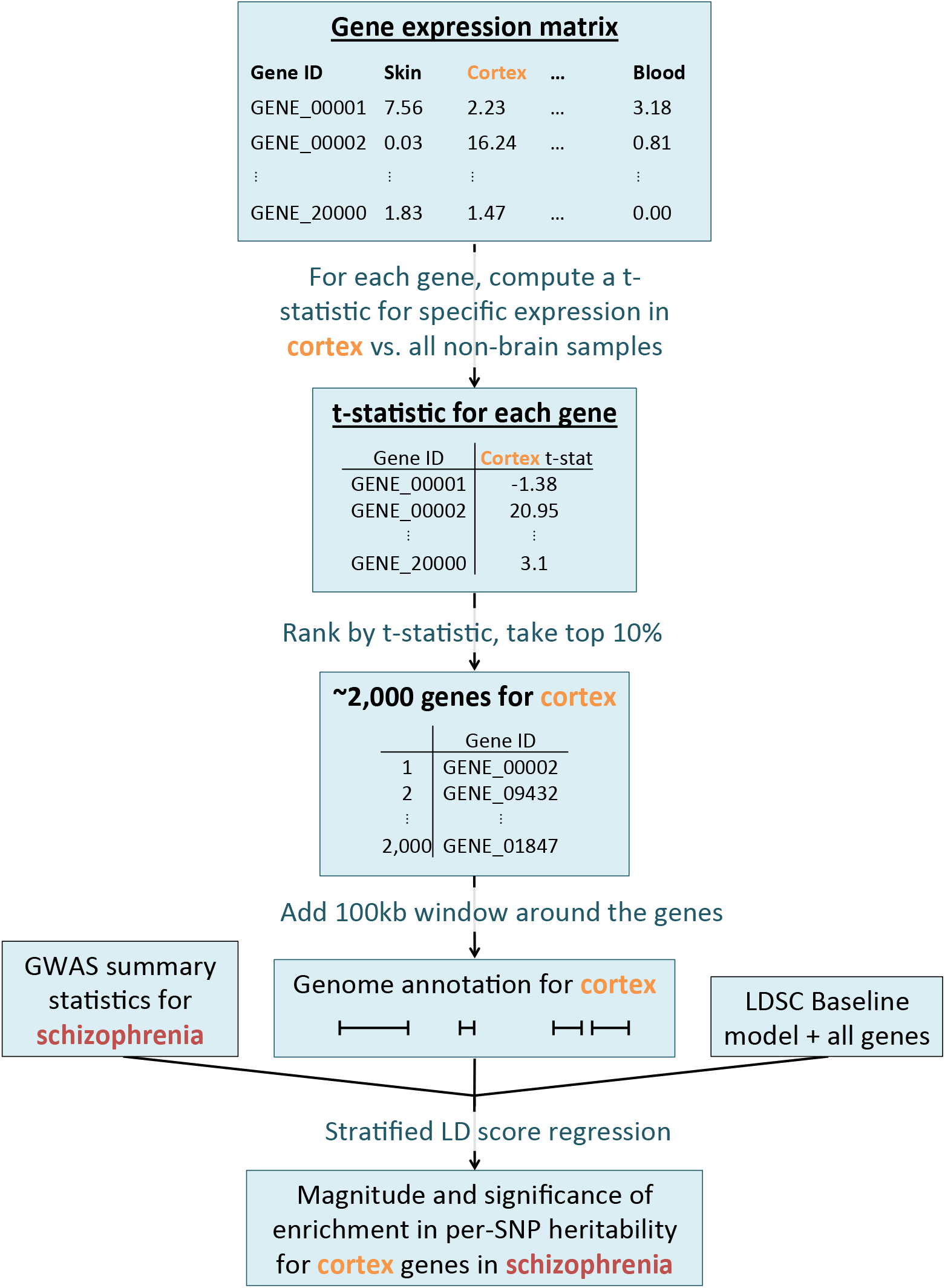
Overview of the approach. For each tissue in our gene expression data set, we compute t-statistics for differential expression for each gene. We then rank genes by t-statistic, take the top 10% of genes, and add a 100kb window to get a genome annotation. We use stratified LD score regression^7^ to test whether this annotation is significantly enriched for per-SNP heritability, conditional on the baseline model^7^ and the set of all genes.

### Analysis of 48 complex traits across multiple tissues

We first analyzed two gene expression data sets. The first data set, from the GTEx consortium v6p^3^, consists of RNA-seq data for 53 tissues, with an average of 161 samples per tissue (**Table S2**, Online Methods). The second data set, which we call the Franke lab data set, is an aggregation of publicly available microarray gene expression data sets comprising 37,427 samples in human, mouse, and rat^16,18^. After removing redundant data, this data set contained 152 tissues, including much better representation of immune tissues and cell types than the GTEx data set (**Table S3**, Online Methods). The gene expression values in the publicly available Franke lab data set already quantify relative expression for a tissue/cell-type rather than absolute expression for a single sample^16,18^, and so we used these values in place of our t-statistics. For visualization purposes, we classified the 205 tissues and cell types in these data sets into nine categories; the classification is described in **Table S2** and **Table S3**. The main goal of this multiple-tissue analysis was to identify system-level enrichments.

We analyzed GWAS summary statistics for 48 diseases and traits with an average sample size of 169,331 (**Table S4**), applying LDSC-SEG for each of the 205 specifically expressed gene annotations in turn. The 48 traits included 13 traits from the UK Biobank^25^ (Online Methods), 16 traits with publicly available GWAS summary statistics^26-36^, and 19 traits from the Brainstorm Consortium^17,37-45^. We excluded the HLA region from all analyses, due to its unusual genetic architecture and pattern of LD. For 34 of the 48 traits, at least one tissue was significant at FDR<5% (Figure 2, Figure S1 and Tables S5 and S6). Several of our results recapitulate known biology: immunological traits exhibit immune cell-type enrichments, psychiatric traits exhibit strong brain enrichment, LDL and triglycerides exhibit liver-specific enrichments, BMI-adjusted waist-hip ratio exhibits adipose enrichment, type 2 diabetes exhibits enrichment in the pancreas, and height exhibits enrichments in a variety of tissues in a pattern similar to previous analyses of this trait^19^. In addition, several of our results validate very recent findings from other genetic analyses: in particular, smoking status, years of education, BMI, and age at menarche show robust brain enrichments that recapitulate results from our previous analysis of genetic data together with chromatin data^7^. Our results were robust to the choice of percent of genes used (10%) and to the size of the window used (100kb) (Figure S2). We assessed correlations in enrichment patterns for pairs of traits (Online Methods), and found large and significant correlations among many brain-related phenotypes, among many immune-related phenotypes, and among a third set of phenotypes including height and blood pressure that tended to have enrichments in the musculosketal/connective, cardiovascular, and other categories (Figure S3).

**Figure 2:**
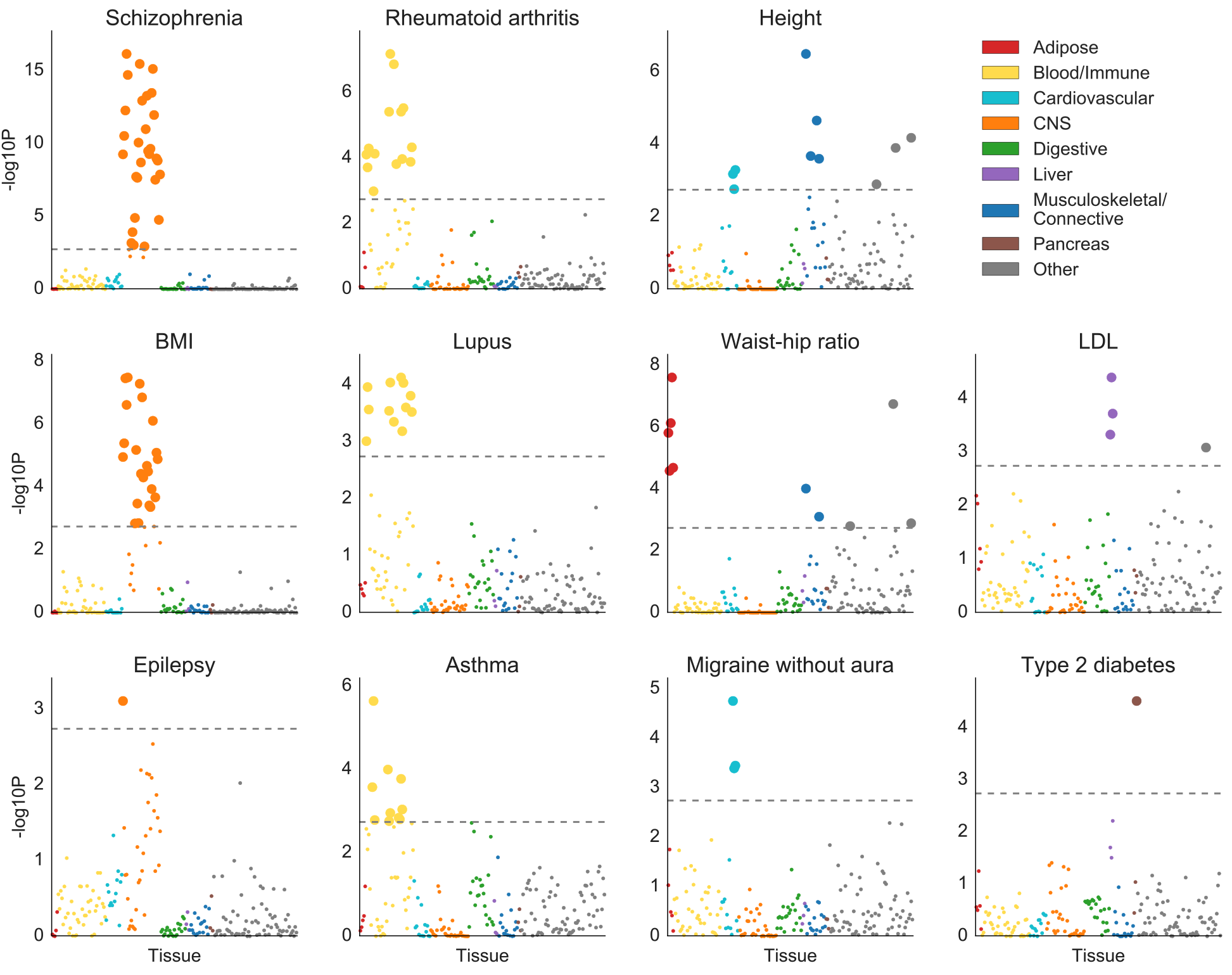
Results of multiple-tissue analysis for selected traits. Results for the remaining traits are displayed in Figure S1. Each point represents a tissue/cell type from either the GTEx data set or the Franke lab data set. Large points pass the FDR<5% cutoff, −log10(P)=2.75. Numerical results are reported in **Table S6.**

Averaging across the most significant tissue for each of these 34 traits, the specifically expressed gene annotation spanned 16% of the genome and explained 36% of SNP-heritability, a 2.3x enrichment (**Table S5**). The sizes of the annotations varied from 11% to 23% of the genome, due to gene size, amount of overlap in the windows around genes, and differences in the number of genes passing QC in the Franke lab data set and the GTEx data. The most significant annotations for each trait explained between 21% and 62% of SNP-heritability, with enrichments varying from 1.4x to 4.7x.

In a data set with many tissues/cell types, related tissues will have highly overlapping gene sets. Because of this, and because we fit each tissue without adjusting for the other tissues analyzed, related tissues often appear enriched as a group. In this analysis, we are focused on identifying system-level enrichments, and so these correlated results do not limit interpretability. The following section similarly focuses on identifying system-level enrichments, while in later sections we focus on differentiating among related tissues/cell types within a system. We note also that the correlation structure among annotations can lead to a distribution of P values that is highly non-uniform, with many P-values close to 0 or 1; this is caused by our one-sided test for enrichment, testing whether the regression coefficient–which represents the change in per-SNP heritability due to a given annotation, beyond what is explained by the set of all genes as well as the baseline model–is positive. The P-values near 0 occur due to correlated annotations with true signal, and the P-values near 1 occur due to annotations without true signal that, conditional on the baseline model, are negatively correlated to annotations with true signal as a consequence of our construction of sets of specifically expressed genes; these annotations thus have negative regression coefficients.

### Validation using independent chromatin data

We analyzed the same 48 diseases and traits using stratified LD score regression^7^ in conjunction with chromatin data from the Roadmap Epigenomics and ENCODE projects^1,2^ (see URLs) instead of gene expression data, with three goals: (1) to validate the results from our analysis of gene expression data using a different type of data from an independent source (2) to identify new enrichments using chromatin data that we did not observe using gene expression data, and (3) to compare enrichments from the two types of data. Using Roadmap data, we constructed 396 cell-type-/tissue-specific annotations from narrow peaks in six chromatin marks—DNase hypersensitivity, H3K27ac, H3K4me3, H3K4me1, H3K9ac, and H3K36me3—each in a subset of a set of 88 primary cell types/tissues. This analysis differed from our previous analysis of chromatin data^7^ in that we used more recently available data on a larger set of chromatin marks, we used peak calls for all marks, and we controlled not only for the union of annotations for each mark, but also for the average of annotations for each mark (Online Methods). We used a subset of the ENCODE data from a subproject called EN-TEx, which includes epigenetic data on a set of tissues that match a subset of the tissues from the GTEx project but are from different donors. Specifically, we used EN-TEx data to construct 93 annotations from peaks for four chromatin marks—H3K27ac, H3K4me3, H3K4me1, and H3K36me3—each in a subset of a set of 27 tissues that were also included in the GTEx data set. We analyzed GWAS summary statistics for the 48 traits, applying stratified LD score regression to each of the 489 tissue-specific chromatin-based annotations in turn.

We considered two types of validation for the results of the multiple-tissue analysis of gene expression described above: validation at the system level and validation at the tissue/cell-type level. For validation at the system level, we classified the top tissue or cell type for each trait with a significant enrichment into one of nine categories (Online Methods), and we considered an enrichment to be validated if a tissue or cell type from the same system passed FDR < 5% for the same phenotype in the chromatin analysis. For validation at the tissue/cell-type level, we only analyzed the 27 tissues present in both GTEx and EN-TEx, and we considered an enrichment of a tissue in GTEx to be validated if any mark in the same tissue in EN-TEx passed FDR < 5% for the same phenotype. The top enrichment from our multi-tissue analysis of gene expression was validated at the system level for 33 out of 34 phenotypes (Figure 3a, **Table S5**). Of these, 29 enrichments had previously been identified in analyses of chromatin data^6,7,37,46-48^ and one had previously been identified in analyses of gene expression data^17^. Out of 20 phenotypes with an enrichment of a tissue or cell type shared between GTEx and EN-TEx that passed FDR<5%, the top enrichment was validated at the tissue/cell-type level for 13 phenotypes (**Table S5**). If we allowed an enrichment of any artery sample in GTEx to be validated by an enrichment of any artery sample in EN-TEx (instead of requiring strict matching of aorta, tibial artery, and coronary artery), the number of validations rose from 13 to 16. Of the four remaining results that were not validated, three were an enrichment in lung for an immunological disease; for all three diseases, the top enrichment in the analysis of gene expression (not restricting to tissues shared between GTEx and EN-TEx) was an immune category from the Franke lab dataset, and the top enrichment in the analysis of chromatin data was an immune category in the Roadmap dataset. We hypothesize that the lung samples analyzed in GTEx may have contained substantial amounts of blood and thus exhibited a gene expression signature reflecting immune activity; this is supported by a GO enrichment analysis of the lung gene set, in which the top three results were related to antigen presentation, immune response, and cytokine-mediated signaling, respectively. In many instances, the analysis of chromatin data detected more enrichments and/or enrichments at higher significance levels than the analysis of gene expression data, though this was not always the case (see Figure S4 and Discussion). We note that in analyses of tissues/cell types for which both gene expression and chromatin data are available, it could be of interest to combine these types of data; we leave this challenge to future work (see Discussion).

**Figure 3:**
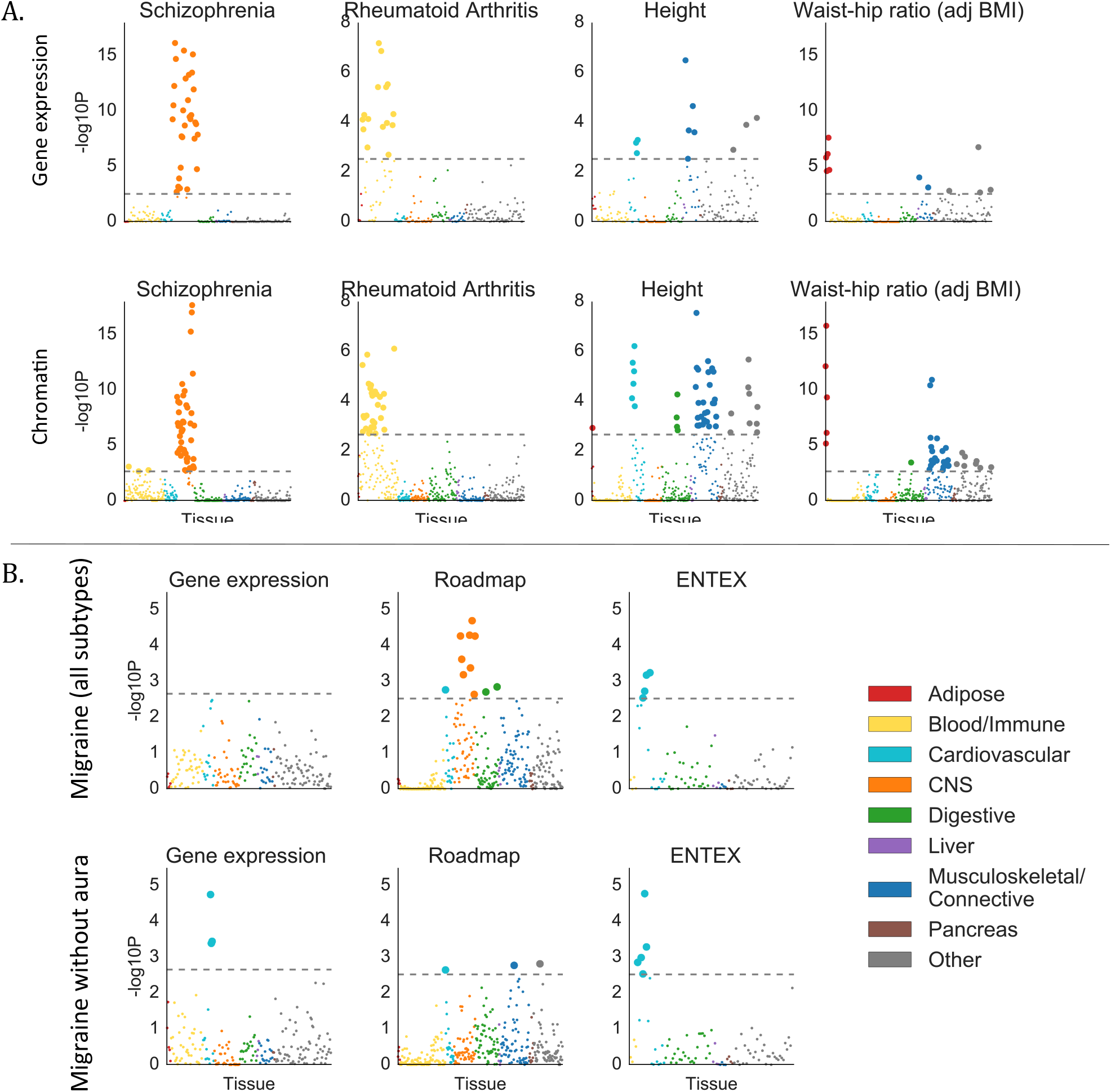
Validation of gene expression results with chromatin data. (A) Examples of validation using chromatin data (bottom) of results from gene expression data (top), for selected traits. Results using chromatin data for all traits are displayed in Figure S5, with numerical results in **Table S7.** For the chromatin results, each point represents a track of peaks for H3K4me3, H3K4me1, H3K9ac, H3K27ac, H3K36me3, or DHS in a single tissue/cell type. (B) Results using gene expression data (including GTEx), Roadmap, and EN-TEx, for migraine (all subtypes) and migraine without aura. For both subfigures, large points pass the FDR<5% cutoff, −logıo(P)=2.85 (chromatin) or −logıo(P)=2.75 (gene expression).

Aggregating all results of the Roadmap and EN-TEx chromatin analyses, at least one tissue was significant at FDR<5% for 44 of the 48 traits (Figure S5 and Tables S5 and S7). The enrichment correlations in this analysis showed a similar pattern to the gene expression analysis above (Figure S6). Averaging across the most significant annotation for each of these 44 traits, the tissue-specific chromatin annotation spanned 3.3% of the genome and explained 43% of the SNP-heritability (**Table S5**). The sizes of the annotation ranged from 8% to 7.8%, and the estimates of enrichment varied from 3.5x to 33x, representing much more variability than for the top annotations in the multiple-tissue gene expression analysis. Because the annotations were much smaller, the estimates of proportion of heritability tended to be much noisier.

Our results for migraine are reported in Figure 3b. We analyzed three migraine GWAS data sets that were recently published^17^: a GWAS of migraine with aura, a GWAS of migraine without aura that had disjoint cases and overlapping controls with the migraine with aura data set, and a migraine (all subtypes) data set that contained the cases from both of these subtypes as well as a large number of additional cases whose subtype was unknown. There is a long-standing scientific debate as to whether migraine has a primarily neurological or vascular basis^49^, and a previous analysis of the migraine (all subtypes) GWAS data together with the GTEx gene expression data reported both cardiovascular and digestive/smooth muscle enrichments^17^. We detected similar cardiovascular enrichments using GTEx gene expression data, although in our analysis the only significant enrichments were for migraine without aura. With EN-TEx, we again saw cardiovascular enrichments, this time for migraine without aura and migraine (all subtypes). Our analysis of Roadmap data, however, yielded qualitatively different results: the strongest enrichment for migraine (all subtypes) was a neurological enrichment that was completely absent in the analysis of migraine without aura, and in the analyses of EN-Tex and gene expression data. The top two annotations were neurospheres and fetal brain, neither of which was present in the gene expression data we analyzed nor in EN-TEx. The correlation in enrichments between migraine (all subtypes) and migraine without aura in the gene expression analysis was estimated to be 0.48 (s.e. 0.15), and in the chromatin data was estimated to be 0.60 (s.e. 13). Our results are consistent with the hypothesis that migraine without aura does indeed have a vascular component, and that another subtype of migraine may have a neurological basis which is sufficiently cell-type specific that the relevant cell types are not represented in either the GTEx or Franke lab data sets. These results highlight the importance of having as many tissues and cell types as possible represented in a multiple-tissue analysis.

A major advantage of gene expression data is that it is available at finer tissue/cell-type resolution within several systems. In the within-system analyses that follow, we investigate these finer patterns of tissue/cell-type specificity.

### Analysis of 12 brain-related traits using fine-scale brain expression data

We identified 12 traits with CNS enrichment at FDR<5% in our gene expression and/or chromatin analyses: schizophrenia, bipolar disorder, Tourette syndrome, epilepsy, generalized epilepsy, ADHD, migraine, depressive symptoms, BMI, smoking status, years of education, and neuroticism. The nervous system has been implicated, either with genetic evidence or non-genetic evidence, for each of these traits^7,27,37,45,49-52^. We first investigated whether some brain regions are enriched over other brain regions for these traits. While the multiple-tissue analysis included annotations for many different brain regions, the gene sets for the different brain regions were often highly overlapping so that for many traits, many brain regions were identified as enriched. For example, nearly every brain region in either the GTEx or Franke lab data was found to be enriched at FDR<5% (Figure 2) in schizophrenia. To differentiate among brain regions, we restricted ourselves to gene expression data only from samples from the brain in the GTEx data. We computed t-statistics within the brain-only data set; e.g. we computed t-statistics for cortex vs. other brain regions instead of cortex vs. other tissues in GTEx, and we used these new t-statistics to construct and test gene sets as in the multiple-tissue analysis. Individual-level data was not available for the Franke lab data set, and thus we could not compute within-brain t-statistics for this data set.

An alternative approach would be to undertake a joint analysis of the original 13 annotations from the multiple-tissue analysis. However, joint analysis of 13 highly correlated annotations is likely to be underpowered, while re-computing t-statistics within the brain allows us to construct new annotations with lower correlations (Figure S7), increasing our power. Moreover, differential expression within the brain may allow us to isolate signals from cell types or processes that are unique to a single brain region, separately from the cell types or processes that are unique to the brain but shared among brain regions. Thus, we use differential expression within the brain, rather than joint analysis of the original annotations, to differentiate among brain regions.

The results of our analysis comparing brain regions are displayed in Figure 4a and **Table S8a**. We identified significant enrichments in the cortex relative to other brain regions at FDR<5% for bipolar disorder, schizophrenia, depressive symptoms, and BMI, and in the striatum for migraine. These enrichments are consistent with our understanding of the biology of these traits^53-56^, but to our knowledge have not previously been reported in any integrative analysis using genetic data. We also identified enrichments in cerebellum for bipolar disorder, years of education, and BMI. However, we caution that differential gene expression in samples from different brain regions can reflect the cell type composition of these brain regions as well as their function. In particular, the cerebellum is known to have a very high concentration of neurons^57^, and thus cerebellar enrichments could indicate either that the cerebellum is a region that is important in disease etiology, or that neurons are an important cell type. While many pairs of the phenotypes had high estimated enrichment correlations in this analysis, migraine tended to have low enrichment correlations with other phenotypes (Figure S8); for example, the estimated enrichment correlation between migraine and schizophrenia was 0.06 (s.e.=0.30) while the estimated enrichment correlation between bipolar disorder and schizophrenia was 0.96 (s.e.=0.05).

**Figure 4:**
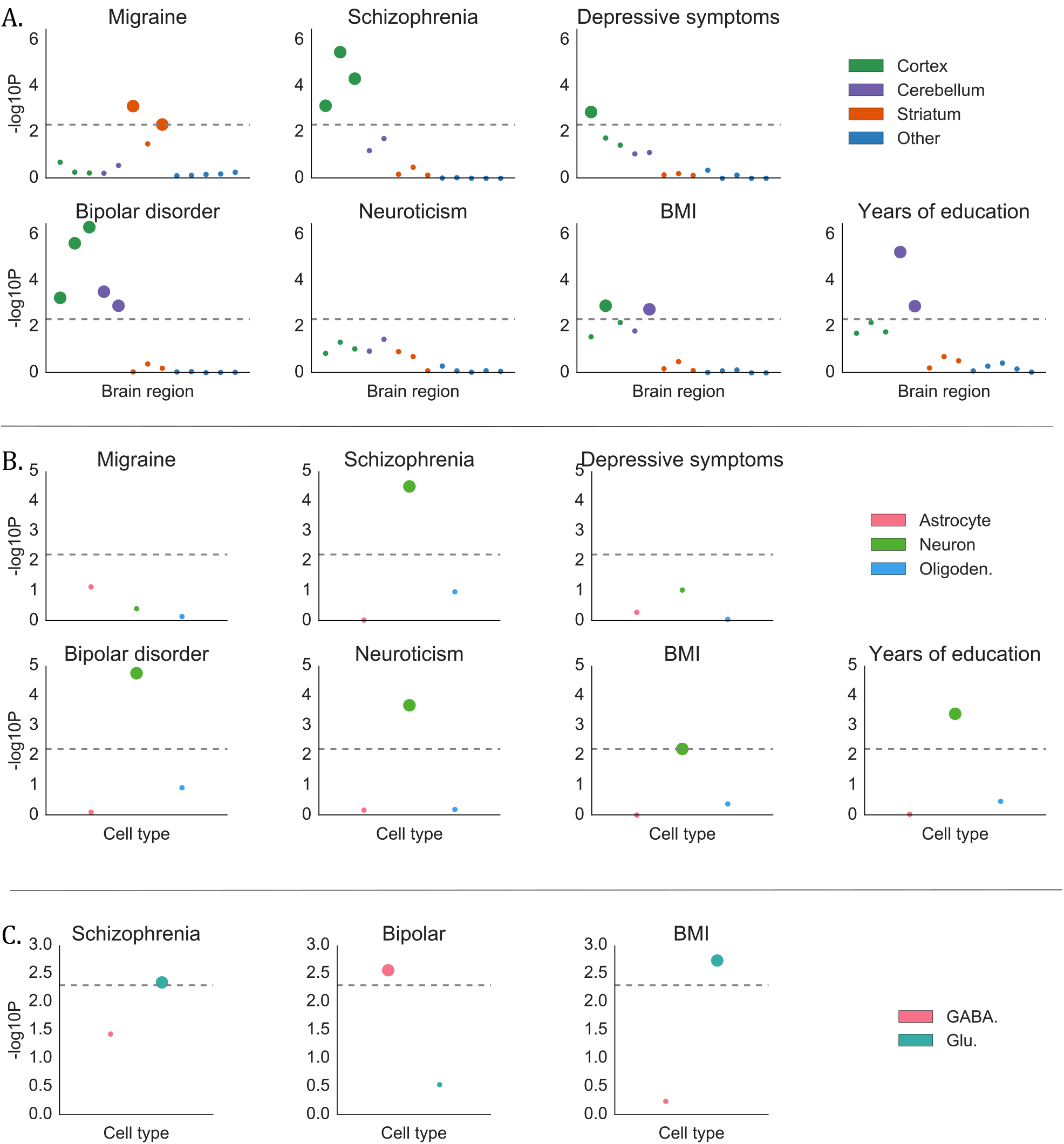
Results of the brain analysis for selected traits, including all significant results. Numerical results for all traits are reported in Table 8.(A) Results from within-brain analysis of 13 brain regions in GTEx, classified into four groups, for seven of 12 brain-related traits. Large points passed the FDR<5% cutoff, −logıo(P)=2.34. (B) Results from the data of Cahoy et al. on three brain cell types for seven of 12 brain-related traits. Large points passed the FDR<5% cutoff, −logıo(P)=2.22. (C) Results from PyschENCODE data on two neuronal subtypes for three of five neuron-related traits. Large points passed the Bonferroni significance threshold in this analysis, −logıo(P)=2.06.

To address the question of the relative importance of brain cell types, as opposed to brain regions, we analyzed the same set of traits using a publicly available data set of specifically expressed genes identified from different brain cell types purified from mouse forebrain^21^. The authors of this data set made lists of specifically expressed genes for each of the three brain cell types available, and these lists were all approximately the same size as the sets of specifically expressed genes in our previous analyses. We created annotations from these lists in the same way that we created annotations from the lists of top 10% expressed genes. The results of this analysis are displayed in Figure 4b and **Table S8b**. We identified neuronal enrichments at FDR<5% for five traits: bipolar disorder, schizophrenia, years of education, BMI, and neuroticism. The other cell types did not exhibit significant enrichment for any of the 13 brain-related traits. The enrichment of neurons for all three of the traits with enrichment in cerebellum in the brain-region analysis supports the hypothesis that analyses of brain regions may be confounded by cell-type composition.

To more precisely characterize the neuronal enrichments, we analyzed the five traits with neuronal enrichment at FDR<5% using t-statistics computed by the PsychENCODE consortium^22^ on differential expression in glutamatergic (excitatory) vs. GABAergic (inhibitory) neurons. The results are displayed in Figure 4c and **Table S8c**; we used Bonferroni correction in this analysis, as we were testing only 5x2=10 hypotheses. For bipolar disorder, genes that are specifically expressed in GABAergic neurons exhibited heritability enrichment, while genes specific to glutamatergic neurons did not. This result supports the theory that pathology in GABAergic neurons can contribute causally to risk for bipolar disorder^58,59^. For BMI and schizophrenia, on the other hand, we found an enrichment in glutamatergic neurons but not in GABAergic neurons.

We were unable to validate the results of these analyses using independent chromatin data. For the two analyses of brain cell types, this was because we were not aware of any available data sets with analogous chromatin data. For the analysis of brain regions, this was because the chromatin annotations that we analyzed were highly correlated across different brain regions and thus some phenotypes were enriched in nearly every brain region; we did not consider these non-specific enrichments to be a meaningful validation of our region-specific results using gene expression data.

### Analysis of 25 immune-related traits using immune cell expression data

We identified 25 traits with immune enrichment at FDR<5% in our gene expression and/or chromatin analyses. This includes many immunological disorders: celiac disease, Crohn’s disease, inflammatory bowel disease, lupus, primary biliary cirrhosis, rheumatoid arthritis, type 1 diabetes, ulcerative colitis, asthma, eczema, and multiple sclerosis. It also includes Alzheimer’s and Parkinson’s diseases, which are neurodegenerative diseases with an immune component previously identified from genetics^60,61^, as well as several brain-related traits–ADHD, anorexia nervosa, bipolar disorder, schizophrenia, Tourette syndrome, and neuroticism–and HDL, LDL, triglycerides, diastolic and systolic blood pressure, hypertension, and BMI. Several of the brain-related traits have been previously suggested to have an immune component^45,62,63^; HDL, LDL, and triglycerides have been linked to immune activation^64-67^; immune cells are causally involved in blood pressure and hypertension^68^; and obesity, in addition to contributing to inflammation^69^, can also be induced in mice through alterations of the immune system^70^. We investigated cell-type-specific enrichments for these traits in 292 immune cell types using gene expression data from the ImmGen project^23^, which contains microarray data on these cell types from mice. This data set contains data for many immune cell types that are not available in the multiple-tissue analysis, and because we compute t-statistics within the data set–i.e., each immune cell vs. all other immune cells–the gene sets are less overlapping than those of immune cell types in the multiple-tissue analysis.

We identified enrichments at FDR<5% for 16 traits. Results are displayed in Figure 5, Figure S9 and Tables S9 and S10, and reveal highly trait-specific patterns of enrichment. For primary biliary cirrhosis, the largest and most significant enrichment was in B cells, consistent with literature on the importance of B cells for this trait^71,72^. Alzheimer’s disease exhibits enrichment in myeloid cells, as seen previously from genetics^73,74^. Asthma and eczema both exhibited enrichment in T and NKT cells; several subclasses of T cells have been shown to be important in asthma,^75^ and a previous study using chromatin data found an enrichment in T cells for asthma but not in other immune cell types^6^. Rheumatoid arthritis, Crohn’s disease, inflammatory bowel disease, and multiple sclerosis all exhibited enrichments in a variety of cell types, consistent with complex etiologies for these diseases that involve many different immune cell types^76-78^. Schizophrenia and bipolar disorder both exhibited an enrichment in T cells. Patients with bipolar disorder have been shown to have a reduction in certain types of T cells, but have equal levels of B cells, NK cells, and monocytes compared to controls^79^. T cell levels have been shown to vary between schizophrenia cases and controls, but existing literature is not consistent in its description of the direction of effect^80^. Note that our analysis excludes the HLA region; a previous analysis of the HLA region for schizophrenia implicated the complement system through its role in synaptic pruning, a signal that is distinct from the signal we observe here^81^. Finally, we identified an enrichment in stromal cells for both diastolic and systolic blood pressure. For each of these two traits, we identified enrichments in the musculoskeletal/connective category in the multiple-tissue analysis that were stronger than the immune enrichments in that analysis, and thus we hypothesize that the enrichment in stromal cells is not providing better resolution on the immune enrichment but instead reflects the more general importance of connective tissue. In enrichment correlation analyses, schizophrenia and bipolar disorder clustered with immunological diseases, while metabolic traits, neurological diseases, and other psychiatric diseases did not (Figure S10).

**Figure 5:**
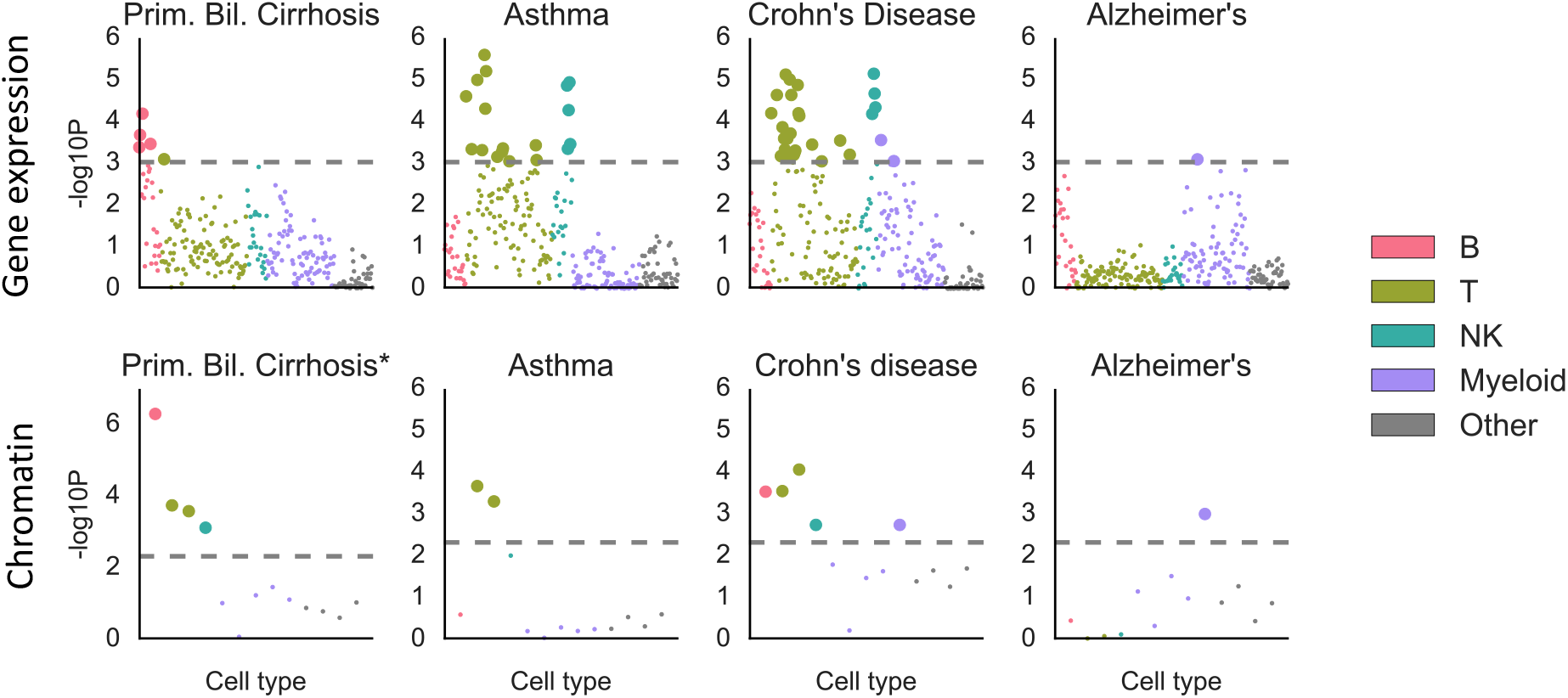
Results of the analysis of ImmGen gene expression data (top) and hematopoiesis ATAC-seq data (bottom) for selected traits. Results for the remaining traits are displayed in Figure S9. Large points passed the FDR<5% cutoff, −log1ø(P)=3.03 (Gene expression) or −log10(P)=2.32 (Chromatin). Numerical results are reported in **Table S10**.

To validate these results, we analyzed ATAC-seq (chromatin) data from 13 cell types spanning the hematopoietic hierarchy in humans^82^. The 13 cell types did not allow us to validate at very high resolution; instead, we classified all cell types from ImmGen and from the hematopoiesis data set into five categories: B cells, T cells, NK cells, myeloid cells, and other cells. There were no stromal cells in the hematopoiesis data set and it was not possible to validate the enrichments for diastolic and systolic blood pressure; this left us with 14 phenotypes with an enrichment at FDR<5% in the ImmGen analysis where the top result fell into one of the first four categories (excluding “Other”). We considered one of these 14 results to be validated if any cell type in the same category from the hematopoiesis data set passed FDR < 5%. Of the 14 phenotypes, the top result was validated for 10 phenotypes (**Table S9**). The only immunological disease whose result was not validated was lupus; the top result for lupus in the ImmGen analysis was a myeloid cell type, while the largest and most significant enrichment in the hematopoiesis data set was a B cell enrichment, consistent with other genetic studies of this trait^14^. The other three phenotypes whose top results did not replicate were schizophrenia and bipolar disorder (T helper cells) and neuroticism (mast cells).

## DISCUSSION

We have shown that applying stratified LD score regression to sets of specifically expressed genes identifies disease-relevant tissues and cell types. Our approach, LDSC-SEG, allows us to take advantage of the large amount of gene expression data available–including finegrained data for which we do not currently have a comparable chromatin counterpart–to ask questions ranging in resolution from whether a trait is brain-related to whether excitatory or inhibitory neurons are more important for disease etiology. We identified many significant enrichments that confirm or extend our current understanding of biology, including an enrichment of striatum for migraine, enrichment of inhibitory neurons for bipolar disorder and excitatory neurons for schizophrenia and BMI, and an enrichment of T cells for Asthma. These results improve our understanding of these diseases, and highlight the power of GWAS as a source of biological insight. Our results may also be useful for choosing the relevant tissue or cell type for in-vitro experiments to further elucidate molecular mechanisms underlying genome-wide significant loci identified in genome-wide association studies. Additionally, cell type-specific annotations have been shown to be useful for fine-mapping^11^, but to our knowledge, all existing methods for functionally informed fine-mapping only examine enrichments inferred from genome-wide significant loci; our work makes us optimistic that enrichments inferred from genome-wide data will further improve fine-mapping resolution and gene prioritization, if appropriate methods can be developed.

There are several key differences between LDSC-SEG, which relies on gene expression data without genotypes or eQTLs, and approaches that require eQTL data^3,13^. First, our approach can be applied to expression data sets such as the Franke lab data set, the Cahoy data set, the PsychENCODE data set, and the ImmGen data set that do not have genotypes or eQTLs available (Table 1). Second, to our knowledge, no method based on eQTLs has been shown to consistently identify system-level enrichments such as brain enrichments for psychiatric traits and immune enrichment for immunological traits, as we do here^3,13,83,84^. For example, a recent study^84^ tested 30 phenotypes for tissue-specific enrichment in 44 tissues from GTEx using the TWAS approach^85^ but concluded that their results “did not suggest tissue-specific enrichment at the current sample sizes.” We share their hypothesis that this is because eQTLs are often shared across tissues even when overall expression levels are very different. Third, methods based on eQTLs require gene expression sample sizes that are large enough to detect eQTLs. In an analysis of data from the GTEx project, we determined that we could identify strong enrichments such as brain enrichment for schizophrenia with just one brain sample, though subtler enrichments had decreasing levels of significance as the gene expression data were down-sampled (Figure S11, Online Methods). Results from our analysis of ImmGen data, which has 2.8 samples per cell type on average, confirm that LDSC-SEG can identify significant enrichments even when the gene expression data has a small number of samples per tissue/cell type, in contrast to eQTL-based methods.

Our polygenic approach also differs from other gene expression-based approaches such as SNPsea^14,15^ and DEPICT^16^, which restrict their analyses to subsets of SNPs that pass a significance threshold. For comparison purposes, we repeated the multiple-tissue analysis using SNPsea and DEPICT. We also repeated the multiple-tissue analysis by analyzing our annotations using MAGMA, a recently developed gene set enrichment method^86^ instead of stratified LD score regression^7^. Results are displayed in Figures S12-S15 (see Online Methods). Many broad patterns were consistent across all approaches: immune enrichment for many immunological diseases, liver enrichment for lipid traits, adipose enrichment for BMI-adjusted waist-hip ratio, and enrichment in several tissues for height and heel T-score. However, there were also several discrepancies. First, SNPsea and DEPICT, the two approaches based on top SNPs, did not identify many of the CNS enrichments for brain-related traits identified by LDSC-SEG and by MAGMA. Second, DEPICT and MAGMA identified more enrichments than LDSC-SEG overall, including some enrichments with unclear relationships to known biology. We hypothesized that LDSC-SEG did not identify some of these enrichments because we jointly model our gene expression-based annotations with the many potential genomic confounders that are included in the baseline model (e.g. exons). We conducted simulations that confirmed that LDSC-SEG is the only one of the approaches tested that is well-powered to identify true enrichments for polygenic traits while avoiding genomic confounding (Figure S16; see Online Methods). We note, however, that MAGMA has an option to include gene-level covariates and that the inclusion of such covariates could ameliorate genomic confounding when using MAGMA; we did not explore that option here.

We cannot conclusively say whether gene expression or chromatin data is preferable when both types of data are available in the same tissues and cell types. Our estimated enrichments were higher for the chromatin-based annotations than for the gene expression-based annotations, but the gene expression-based annotations are larger and have less LD to the rest of the genome. Some chromatin marks tend to be more cell type-specific than overall gene expression, but our specifically expressed gene sets have low correlation across tissues (Figure S17). There were two instances in which we had gene expression and chromatin data on the same set of tissues/cell types, and we compared the P-values in our analyses of these data sets. First, we compared our results from GTEx (gene expression) and EN-TEx (chromatin) for the tissues shared between these two data sets in the multiple-tissue analysis, and we found that the two data sets had comparable distributions of P-values (Figure S4). On the other hand, the hematopoietic data set that we analyzed^82^ had matched ATAC-seq and RNA-seq data, and while our analysis of the ATAC-seq peaks lead to significant enrichments for many traits (Figure 5, **Table S10**), the RNA-seq data set yielded only a single enrichment for a single trait (**Table S11**). This leads us to conclude that the question of which type of data is preferable may depend on complex factors such as which chromatin marks were analyzed, the overall quality of the data set, the sample size with which the specifically expressed genes are called, and how similar the tissues are to each other. When gene expression and chromatin data are available on the same set of tissues or cell types, it may be possible to combine these types of data to improve power. For example, it may be useful to restrict an annotation to tissue-specific chromatin marks near specifically expressed genes, or to combine the P-values from separate analyses of the two types of data. We defer a thorough exploration of this set of possibilities to future work.

Our work is based on the assumption that a tissue or cell type is important for a particular disease if and only if SNPs near genes with high specific expression in that tissue/cell type are enriched for heritability. This assumption leads to several limitations of our approach. First, when analyzing gene expression data from different tissues, cell type composition can confound the analysis, as we demonstrated in our comparison of brain regions; this makes enrichments of organs such as the esophagus or uterus hard to interpret. Second, tissues/cell types with similar gene expression profiles to a causal tissue/cell type will be identified as relevant to disease, just as SNPs in LD with a causal SNP will be identified as associated to disease in a GWAS; thus, significant tissues/cell types should be cautiously interpreted as the “best proxy” for the truly causal tissue/cell type, which may be unobserved. Third, our focus on nearby SNPs prevents us from leveraging signal from regulatory SNPs that act at longer distances. Our approach is also fundamentally limited by the availability of gene expression data and cannot rule out the importance of a given cell type; for example, if the tissue/cell type that is most relevant for a disease occurs in a stage of development or under a stimulus that has not been assayed, then we may not identify enrichments in that tissue/cell type.

Our use of a heritability-based approach has advantages but also leads to some limitations. First, our approach will not detect strong but highly localized signals. Second, power increases only modestly with sample size at very large sample sizes, as the finite size of the genome is a stricter constraint for highly heritable traits at these sample sizes: for example, LD score regression coefficients of baseline model annotations had s.e. that were only 1.29x lower on average in analyses of the full UK Biobank data set (average N=438,682) vs. the interim UK Biobank data set (average N=140,026.) Also, because our approach uses stratified LD score regression, it cannot be applied to custom array data, it requires a sequenced reference panel that matches the population studied in the GWAS, and we cannot rule out bias due to model misspecification^7^. Augmentations to the baseline model^87^ may help ameliorate potential model misspecification, but we leave further investigation of this to future work.

Another limitation of our method is that its results may be difficult to validate. We undertook a type of validation using independent chromatin data, when there was comparable chromatin data available. However, this type of validation involves a number of challenges. First, we often do not have chromatin data in the same tissues and cell types as the gene expression data. Second, it is not clear that we should always expect results to replicate; for example, it is biologically plausible that SNPs near specifically expressed genes in the relevant tissue are enriched, while SNPs in H3K36me3 peaks called in the tissue are not. Third, our gene expression annotations represent relative activity—we select genes with higher expression in the focal tissue compared to other tissues—while the chromatin annotations that we use here represent absolute activity (although relative chromatin annotations are also possible^6,88^). Despite these limitations, replicating an enrichment for a particular system, tissue, or cell type using independent chromatin data can provide a strong validation for gene expression results.

Our power to identify disease-relevant tissues and cell types will improve as large GWAS sample sizes become available for more phenotypes, and as gene expression data is generated in new tissues and cell types. This will help advance our understanding of disease biology and lay the groundwork for future experiments exploring specific variants and mechanisms.

## ACKNOWLEDGEMENTS

We are thankful to Masahiro Kanai, Farhad Hormozdiari, Jacob Ulirsch, Tune Pers, Sam Riesenfeld, Rebecca Herbst, Adrian Veres, and Eran Hodis for helpful comments. This research has been conducted using the UK Biobank Resource (Application Number: 16549). This research was funded by NIH grants R01 MH107649, R01 MH109978 and U01 CA194393. HKF is supported by the Fannie and John Hertz Foundation. The data on neuron types were generated as part of the PsychENCODE Consortium, supported by: U01MH103339, U01MH103365, U01MH103392, U01MH103340, U01MH103346, R01MH105472, R01MH094714, R01MH105898, R21MH102791, R21MH105881, R21MH103877, and P50MH106934 awarded to: Schahram Akbarian (Icahn School of Medicine at Mount Sinai), Gregory Crawford (Duke), Stella Dracheva (Icahn School of Medicine at Mount Sinai), Peggy Farnham (USC), Mark Gerstein (Yale), Daniel Geschwind (UCLA), Thomas M. Hyde (LIBD), Andrew Jaffe (LIBD), James A. Knowles (USC), Chunyu Liu (UIC), Dalila Pinto (Icahn School of Medicine at Mount Sinai), Nenad Sestan (Yale), Pamela Sklar (Icahn School of Medicine at Mount Sinai), Matthew State (UCSF), Patrick Sullivan (UNC), Flora Vaccarino (Yale), Sherman Weissman (Yale), Kevin White (UChicago) and Peter Zandi (JHU).

## URLs

- LDSC software, including LDSC-SEG: https://github.com/bulik/ldsc.
- Gene sets and LD scores from this paper: https://data.broadinstitute.org/alkesgroup/LDSCORE/.
- GTEx: http://www.gtexportal.org.
- Franke lab data: https://data.broadinstitute.org/mpg/depict/depict_download/ tissue_expression.
- Cahoy et al. data: http://jneurosci.org/content/suppl/2008/01/03/ 28.1.264.DC1, see Tables S4-S6.
- PsychENCODE: https://www.synapse.org/#Synapse:syn4921369/wiki/235539.
- ImmGen, https://www.immgen.org/.
- Roadmap Epigenomics: http://www.roadmapepigenomics.org.
- GERA data set (database of Genotypes and Phenotypes (dbGaP), phs000674.v1.p1): http://www-ncbi-nlm-nih-gov.libproxy.mit.edu/projects/gap/cgi-bin/study.cgi?study_id=phs000674.v1.p1.
- PLINK: https://www.cog-genomics.org/plink2.
- makegenes.sh: https://github.com/freeseek/gwaspipeline

## ONLINE METHODS

### Computing t-statistics

When computing the t-statistic of each gene for a focal tissue, we excluded all samples from the same tissue category (see “Tissue categories and covariates” below). For example, when computing the t-statistic of specific expression for each gene in cortex using GTEx data, we compared expression in cortex samples to expression in all other samples, excluding other brain regions. We chose to exclude other brain regions because we wanted to include genes that are more highly expressed in brain tissues than in non-brain tissues, even if they are not specific to cortex within the brain. This procedure results in a higher correlation among the t-statistics for the different brain regions; in a separate analysis, we compute within-brain t-statistics to disentangle this signal.

Thus, for a focal tissue (e.g., cortex) in a larger tissue category (e.g., brain), we computed the t-statistic for gene g as follows. We first constructed a design matrix *X* where each row corresponds to a sample either in cortex or outside of the brain. The first column of *X* has a 1 for every cortex sample and a -1 for every non-brain sample. The remaining columns are an intercept and covariates (see “Tissue categories and covariates” below). The outcome *Y* in our model is expression. We fit this model via ordinary least squares, and compute a t-statistic for the first explanatory variable in the standard way:

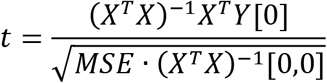

where MSE is the mean squared error of the fitted model; i.e.,

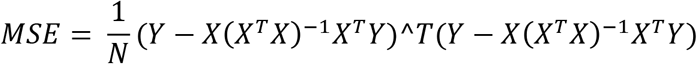

where *N* is the number of rows in X. This gives us a t-statistic for each gene for the focal tissue. We then select the top 10% of genes, add a 100kb window around their transcribed regions, and apply stratified LD score regression to the resulting genome annotations as described below.

### Modifications of our approach

For some analyses, we modified our approach to constructing sets of specifically expressed genes to better take advantage of the data available.

- *Franke lab data set.* The values in the publicly available matrix are not a quantification of expression intensity, but rather a quantification of differential expression relative to other tissues in this data set^16^,^18^. Thus, it was not appropriate to compute t-statistics in this data set. We used the original values in place of our t-statistics, then proceeded as described in Figure 1.
- *Cahoy data set.* The data set of Cahoy et al. had available sets of specifically expressed genes for the three cell types that each had between 1,700 and 2,100 genes. We took these to be the gene sets for the three cell types, then proceeded as in the standard approach, adding a 100kb window and applying stratified LD score regression.
- *PsychENCODE data set.* The PsychENCODE data set had available t-statistics for GABAergic neurons vs. Glutamatergic neurons. We used these t-statistics, rather than computing our own.

For the other data sets we analyzed (GTEx, GTEx brain regions, ImmGen), we used the approach described in Figure 1. We view it as an advantage of our method that it can be flexibly adapted to many different types of data.

### Tissue categories and covariates.

- For the multiple-tissue GTEx analysis, we used the “SMTS” variable (“Tissue Type, area from which the tissue sample was taken”) to define the tissue categories (**Table S2**). We used age and sex as covariates.
- For the analysis of GTEx brain regions, we set each tissue to be its own category, and we used age and sex as covariates.
- For the ImmGen analysis, we defined tissue categories using the classification on the main page of immgen.org of cell types into categories: B cells, gamma delta T cells, alpha beta T cells, innate lymphocytes, myeloid cells, stromal cells, and stem cells (**Table S10**). The classification at immgen.org also has a “T cell activation” category that we collapsed into the alpha beta T cell category because it had data on alpha beta T cells at different stages of activation. We did not have any covariates.
- For the Franke lab data set, Cahoy data set, and PsychENCODE data set, we did not compute t-statistics and so we did not have tissue categories or covariates (see “Modifications of our approach” above).

### Choice of parameters

Our approach includes two parameters: the proportion of genes selected, which we set to 10%, and the window size around each gene, which we set to 100kb. To choose these two parameters, we ran the approach with six different parameter settings ({2%, 5%, 10% of genes} x {20kb, 100kb windows}) on two diseases— schizophrenia and rheumatoid arthritis—and two corresponding GTEx tissues—brain (all brain regions) and blood (LCLs and whole blood)—which are widely known to be disease-relevant tissues. We determined that of the parameter settings we tested, 10% of genes and 100kb produced the most significant P-values for identifying brain enrichment for schizophrenia and blood enrichment for rheumatoid arthritis, so we used these parameters for the remaining analyses.

### Application of stratified LD score regression

Stratified LD score regression^7^ is a method for partitioning heritability. Given (potentially overlapping) genomic annotations *C_1_,*…, *C_ĸ_*,one of which is the category of all SNPs, we model the causal effect of SNP *j* on phenotype *Y* as drawn from a distribution with mean 0 and variance

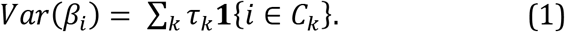

(If the genomic annotations are real-valued rather than subsets of SNPs, we can replace 1{í e *C*_k_} with any other function of the SNP indices^87^.) We then model the phenotype *Y* as depending linearly on genotype: *Y = X ■ ß + e,* where *X* is a vector of SNP values for an individual, and each SNP has been standardized to mean 0 and variance 1 in the population. Because each SNP is standardized, and because *β_t_* has mean zero, we can call *Var*(ß_*i*_) the per-SNP heritability of SNP i. (Note that here, because we model *β* as random, our definition of heritability is different from definitions of heritability in which *β* is fixed, and so we are estimating a fundamentally different quantity than some other methods^89^.)

Under this model, the expected marginal chi-square association statistic for SNP *i* reflects the causal contributions not only of SNP *i* but of SNPs in LD with SNP i. Specifically,

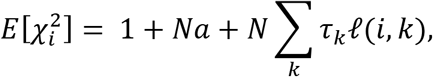

where *N* is the GWAS sample size, *a* is a constant that reflects population structure and other sources of confounding,^90^ and í(*i, k*) is the LD score of SNP *i* to category *C_k_*, defined as í(i,k) = Σ;*r*^2^(í,*j* 1 *{j* E *C_k_*}, where r^2^(*i,j*) is the squared correlation between SNPs *i* and *j* in the population. To estimate the τ_k_, we first estimate •£(í, fc) from a reference panel, and we then perform weighted regression 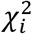 on *N ■* £(*i, k*), using a jackknife over blocks of SNPs to estimate standard errors.

The regression coefficient τ_k_ quantifies the importance of annotation *C_k_*, correcting for all other annotations in the model; *τ_k_* will equal zero if *C_k_* is not enriched, will be negative if belonging to *C_k_* decreases per-SNP heritability accounting for all other annotations included, and will be positive if belonging to *C_k_* increases per-SNP heritability, accounting for all other factors. Thus, as in our previous cell-type-specific anlaysis^7^, we compute P-values that test whether *τ_k_* is positive. When reporting quantitative results, we normalize the coefficient *τ_k_* by our estimate of the mean per-SNP heritability Σ_*i*_ *Var*(ß_*i*_)/*M* to make it comparable across phenotypes. The normalized coefficient can be interpreted as the proportion by which the per-SNP heritability of an average SNP would increase if *τ_k_* were added to it. In addition, it is possible to estimate the total heritability, defined as Σ_*i*_ *Var*(ßļ), as well as the heritability in category *C*_k_, defined as Σ_*i*∈*C*_*k*__ *Var(ß*_i_), by plugging estimates of *τ_k_* into Equation (1), and to compare the proportion of heritability, Σ_*i*∈*C*_*k*__ *Var*(ß_*i*_)/Σ*i V¤·r(ßi*), to the proportion of SNPs, *C_k_*|/*M,* where *M* is the total number of SNPs^7^.

We analyzed autosomes only and excluded the HLA from all analyses. In each analysis, we jointly fit the following annotations:

1. The annotation created for our focal tissue by adding 100kb windows around the top 10% of genes ranked by t-statistic.
2. An identical annotation created for all genes included in the gene expression data set being analyzed.
3. The baseline model with 52 functional categories, described previously^7^ and listed in **Table S1.**

### Gene expression data: quality control and normalization.

- *GTEx data set.* We downloaded the RNA-seq read counts from GTEx v6p (see URLs), removed genes for which fewer than 4 samples had at least one read count per million, removed samples for which fewer than 100 genes had at least one read count per million, and applied TPM normalization^91^. We used the “SMTSD” variable (“Tissue Type, more specific detail of tissue type”) to define our tissues (**Table S2**).
- *Franke lab data set.* We downloaded the publicly available gene expression data from the DEPICT website (see URLs). We determined that several pairs of tissues had values that were correlated at r^2^>0.99, including several that had r^2^=1. We pruned our data so that no two tissues had r^2^>0.99. Most of the closely correlated pairs were also biologically closely related so that the interpretation did not depend on which tissue we chose to keep (e.g., plasma and plasma cells, joint and joint capsule). For pairs of tissues where one tissue was more specific than the second, we kept the more specific pair (e.g., nose vs. nasal mucosa, quadriceps muscle vs. skeletal muscle). There were two clusters of highly correlated tissues for which we decided to remove the entire cluster, not keeping any of the tissues, because these clusters had very strong but biologically implausible correlations. The first such cluster was made up of eyelids, conjunctiva, anterior eye segment, tarsal bones, foot bones, and bones of the lower extremity. The second such cluster was made up of connective tissue, bone and bones, skeleton, and bone marrow. After pruning, this data set contained 152 tissues, listed in **Table S3**.
- *Cahoy et al. data set.* We downloaded sets of specifically expressed genes for each of the three cell types (see URLs). To obtain a list of all genes, we also downloaded a list of all genes that passed quality control in their analysis (**Table S3b** of Cahoy et al.). We mapped from mouse to human genes using orthologs from ENSEMBL (see URLs).
- *PsychENCODE data set.* We used the t-statistics released by the PsychENCODE consortium for differential expression in GABAergic vs. Glutamatergic neurons^22^. These t-statistics were computed using limma^92^.
- *ImmGen data set.* We downloaded publicly available gene expression data from the ImmGen Consortium (see URLs). We used both Phase 1 (GSE15907) and Phase 2 (GSE37448) data. The data on GEO were on an exponential scale, so we log transformed the data and mapped to human genes using ENSEMBL orthologs. We tested each of the 297 cell types.

We modified the makegenes.sh script^93^ (see URLs) for some of our data processing.

### UK Biobank data

We analyzed data from the full N=500K UK Biobank release^25^ for 13 traits (P.R. Loh et al., unpublished data). The summary statistics were generated using BOLT-LMM v2.3, an unpublished extension of BOLT-LMM^94^.

### Enrichment correlation

For a pair of phenotypes and a set of tissues/cell types, we defined the enrichment correlation to be the correlation between the regression coefficients corresponding to each tissue/cell type. We estimated the enrichment correlation by correlating the estimates of the regression coefficients, and we quantified uncertainty via block jackknife over 200 sets of consecutive SNPs. We note that when the number of tissues/cell types included is small, the true underlying enrichment correlation may be large even though there is no relationship between the two phenotypes, so we only estimate enrichment correlations when there are at least 10 tissues or cell types.

### Chromatin analysis

We downloaded narrow peaks from the Roadmap Epigenomics consortium for DNase hypersensitivity and five activating histone marks: H3K27ac, H3K4me3, H3K4me1, H3K9ac, and H3K36me3 (see URLs). Each of these six features was present in a subset of the 88 primary cell types/tissues, for a total of 397 cell-type-/tissue-specific annotations. For each of these annotations, we tested for enrichment by adding the annotation to the baseline model (see **Table S1**), together with the union of cell-type-specific annotations within each mark and the average of cell-type-specific annotations within each mark. A positive regression coefficient for a tissue-/cell-type-specific annotation represents a positive contribution of the annotation to per-SNP heritability, conditional on the other annotations. We again computed a P-value to test whether the regression coefficient was positive.

We also analyzed peaks called using Homer from EN-Tex, a subgroup of the ENCODE project, for four activating histone marks: H3K27ac, H3K4m3, H3K4me1, and H3K36me3. Each of these four marks was present in a subset of a set of 27 tissues matching tissues from the GTEx consortium. We repeated the same analysis as described above for the Roadmap dataset, computing the union and average of cell type-specific annotations within each mark, adding them to the baseline model, and then testing for enrichment by adding one annotation at a time to the regression.

### Classification of tissues/cell types for system-level validation of the results of the multiple-tissue analysis of gene expression

We classified the top tissue or cell type for each trait with a significant enrichment into one of the eight systems (excluding “Other”) in the Figure 2 legend. There were three phenotypes whose top tissue fell in the “Other” category; two of these we classified into a new “Reproductive” category. The last one, serous membrane, did not have any comparable tissues in our chromatin data and so we instead attempted to replicate the second most significant result for that phenotype.

Our analysis of chromatin in this work differs from our previous analysis of chromatin data^7^ in three ways. First, we use a larger range of marks and tissues/cell types: every track available from the Roadmap Epigenomics website (see URLs) for any of six activating marks, H3K27ac, H3K4me1, H3K4me3, H3K9ac, H3K36me3, and DHS, in any of the 88 primary tissues and cell types available, for a total of 397 annotations. Second, we used narrow peaks from Roadmap for all of the marks. Previously, we analyzed H3K27ac data from one source^6^ and H3K4me1, H3K4me3, and H3K9ac data from another source^5^,^12^; now that there is a single standard source with uniformly processed data for all marks of interest, we have switched to using this data. Finally, we controlled more strictly for confounders by including the average across cell types of the cell-type-specific annotations for a given mark as an annotation in the model, so that annotations that tend to fall in areas that are more active overall are not falsely interpreted as cell-type-specific signal.

### Number of gene expression samples needed

Because the GTEx consortium data set included tens of samples for many of the tissues, we were able to assess how sensitive our results were to the sample size of the gene expression data set used to construct the gene sets. To do this, we repeatedly sub-sampled our data set to a variety of sample sizes, each time re-creating gene sets using the smaller sub-sampled data set. We chose two results to re-analyze in this way. First, we re-analyzed cortex enrichment for schizophrenia, in which cortex was compared to all non-brain samples and was highly significant (Figure 2). This result was very robust: the enrichment was highly significant in all of our downsampled data sets, even with only a single cortex sample (Figure S11a). We then assessed enrichment for schizophrenia in the within-brain analysis, in which cortex was compared to all other brain regions and was moderately significant (Figure 4a). In this analysis, sample size was more important, and while there was high variance in z-score among random samples at a given sample size, there was a clear trend that increasing the sample size increases the significance of the enrichment on average (Figure S11b). In conclusion, these analyses provide evidence that sample size can be important when the enrichment being identified is near the border of significance, but that our method is well-powered to detect strong signals even with a single sample in the tissue of interest.

### Comparison to existing methods: real phenotypes

To our knowledge, SNPsea^14-15^ is the only existing method that takes as input GWAS summary statistics, together with a matrix of gene expression values, and identifies enriched tissues and cell types. SNPsea leverages only genome-wide significant SNPs, rather than all SNPs, a notable difference from our approach. We ran SNPsea on the summary statistics and gene expression data analyzed in our multiple-tissue analysis; results are displayed in Figure S12. We found that SNPsea identified biological plausible enrichments at high levels of significance for traits such as LDL for which a large proportion of SNP-heritability lies in genome-wide significant loci, but that it was not well-powered for more polygenic traits; for example, it found zero tissues with FDR < 5% for bipolar disorder, while our approach found many brain regions to be enriched at P-values as low as 2e-12 (Figure S1). The lack of power of SNPsea on more polygenic traits is unsurprising, as SNPsea leverages only genome-wide significant loci.

The DEPICT software^16^ includes a method for identifying disease-relevant tissues and cell types from GWAS summary statistics and gene expression data. However, this method takes as input only the GWAS summary statistics and not gene expression data; the method is designed to be run only with the Franke lab data set^16^,^18^, which is built into the software. Thus, DEPICT could not be used to obtain the results in our brain-specific and immune-specific analyses, for which we analyzed data sets that allowed us to differentiate among tissues and cell types within each of these systems. However, DEPICT does perform a multiple-tissue analysis analogous to the Franke lab data set component of our multiple-tissue analysis, and so we ran DEPICT on the set of summary statistics that we analyzed. Like SNPsea, DEPICT is run on a subset of SNPs, but unlike SNPsea, DEPICT documentation recommends that it be run twice, once on SNPs that pass genome-wide significance at 5e-8, and once on SNPs that pass a less stringent threshold of 1e-5; we followed this recommendation, and our results are displayed in Figures S13 and S14. We determined that DEPICT failed to identify some enrichments identified by our analysis of the Franke lab data set, such as brain enrichment for several brain-related traits (epilepsy, Tourette syndrome, neuroticism, and smoking status), but that it identified a large number of enrichments for other traits and tissues that our approach did not find. In simulations described below, we found that DEPICT sometimes reported significant results in the absence of true enrichment.

Our approach, described in Figure 1, has two main steps: constructing a genome annotation from gene expression data, and testing this annotation for enrichment with GWAS summary statistics using stratified LD score regression. We tested whether the success of our approach depended on using stratified LD score regression in the second step by instead analyzing the specifically expressed gene annotations from the first step using MAGMA^86^, a gene set enrichment method that allows inclusion of a window around each gene and leverages all SNPs in the gene set (Figure S15). MAGMA and LDSC-SEG identified many of the same enrichments, but MAGMA identified several enrichments that LDSC-SEG did not. We hypothesized that this may occur because in this analysis, we did not use the option in MAGMA to incorporate gene-level covariates. In simulations described below, we determined that MAGMA can report significant results in the absence of true enrichment due to uncorrected genomic confounding if no covariates are included to ameliorate potential confounding. We leave an exploration of how best to use covariates in MAGMA to account for potential confounding while preserving power for future work.

For comparison purposes, we report LDSC-SEG results for the multiple tissue analysis as a heatmap in Figure S2a, in addition to the scatter plots in Figure 2 and Figure S1.

### Comparison to existing methods: simulated phenotypes

We performed simulations using genotypes from Genetic Epidemiology Research on Aging (GERA) data set^95-97^ with 47,360 individuals and 6,507,309 SNPs with imputation R^2^ > 0.5. We simulated five genetic architectures, where “null” refers to a heritable trait with no tissue-specific enrichment and “causal” refers to a heritable trait with cortex enrichment:

1. (Polygenic null) All SNPs causal, causal SNP effects are drawn independently from a normal distribution with mean zero and constant variance across the genome, with a total heritability of 0.9.
2. (Sparse null) Same as (1), but each SNP has probability 0.001 of being causal.
3. (Exon-enriched null) A SNP is causal if and only if it is in an exon, causal SNP effects are drawn independently from a normal distribution with mean zero and constant variance for all exonic SNPs, with a total heritability of 0.9.
4. (Polygenic causal) We use the annotation corresponding to cortex genes from the multiple-tissue analysis to simulate a true effect. All SNPs are causal, causal SNP effects are drawn independently from a normal distribution with a constant variance within the cortex annotation and constant variance outside of the cortex annotation so that 50% of the total heritability is assigned to the cortex annotation, 50% of the total heritability is distributed uniformly across the genome, and the total heritability is 0.2. We chose a smaller value of heritability in the causal simulations because we wanted to test power to identify true enrichment rather than control of type I error.
5. (Sparse causal) Same as (4), but each SNP has a probability of 0.001 to be causal.

For each genetic architecture, we simulated phenotypes and summary statistics using PLINK^98^ (see URLs) with 100 replicates for each genetic architecture. We then ran the multiple-tissue analysis as described above for every method on each of the simulated data sets, and for each method and each simulated genetic architecture we performed FDR correction within the set of 100 simulated phenotypes. Results are displayed in Figure S16.

Of the five methods tested (LDSC-SEG, SNPsea, DEPICT (1e-5), DEPICT (5e-8), and MAGMA), only LDSC-SEG and SNPsea correctly reported no significant enrichments passing FDR<5% for all 3 null simulations (scenarios 1-3). In particular, DEPICT with a threshold of 1e-5 reported significant enrichments at FDR<5% for all three null simulations (scenarios 1-3), while DEPICT with a threshold of 5e-8 reported significant enrichments at FDR < 5% for the sparse null simulation (scenario 2). MAGMA correctly reported no significant enrichment for the null simulations with no enrichment (scenarios 1-2) but reported a large number of significant enrichments at FDR<5% for the null simulation with enrichment in exons (scenario 3); we note that we ran MAGMA without taking advantage of the option to incorporate gene-level covariates which would likely ameliorate the false positives.

All five methods reported significant cortex enrichments at FDR<5% for the sparse causal simulation (scenario 5), but only MAGMA and LDSC-SEG reported significant cortex enrichments for the polygenic causal simulation (scenario 4). These simulations, together with the analysis of real phenotypes described above, indicate that when MAGMA is run without covariates, only LDSC-SEG and SNPsea control type I error, and that of these two methods, LDSC-SEG is better powered for polygenic traits.

### Data availability

We have also released all genome annotations derived from the publicly available gene expression data that we analyzed at http://data.broadinstitute.org/alkesgroup/LDSCORE/.

### Code availability

Open source software implementing our approach is available at http://www.github.com/bulik/ldsc.

## References

1. The ENCODE Project Consortium. An integrated encyclopedia of DNA elements in the human genome. Nature 489, 57–74 (2012).

2. Kundaje A. et al. Integrative analysis of 111 reference human epigenomes. Nature 518, 317–330 (2015).

3. The GTEx Consortium. The Genotype-Tissue Expression (GTEx) pilot analysis: Multitissue gene regulation in humans. Science 348, 648–660 (2015).

4. Ernst J. etal. Mapping and analysis of chromatin state dynamics in nine human cell types. Nature 473, 43–49 (2011).

5. Trynka G. et al. Chromatin marks identify critical cell types for fine mapping complex trait variants. Nat. Genet. 45, 124–130 (2013).

6. Farh K. K.-H. et al. Genetic and epigenetic fine mapping of causal autoimmune disease variants. Nature 518, 337–343 (2014).

7. Finucane H. K. et al. Partitioning heritability by functional annotation using genome-wide association summary statistics. Nat. Genet. 47, 1228–1235 (2015).

8. Li Y. & Kellis M. Joint Bayesian inference of risk variants and tissue-specific epigenomic enrichments across multiple complex human diseases. Nucleic Acids Res. (2016). doi:10.1093/nar/gkw627

9. Maurano M. T. et al. Systematic Localization of Common Disease-Associated Variation in Regulatory DNA. Science 337, 1190–1195 (2012).

10. Pickrell J. K. Joint Analysis of Functional Genomic Data and Genome-wide Association Studies of 18 Human Traits. Am. J. Hum. Genet. 94, 559–573 (2014).

11. Kichaev G. et al. Integrating Functional Data to Prioritize Causal Variants in Statistical Fine-Mapping Studies. PLoS Genet. 10, e1004722 (2014).

12. Gusev A. et al. Partitioning heritability of regulatory and cell-type-specific variants across 11 common diseases. Am. J. Hum. Genet. 95, 535–552 (2014).

13. Ongen H. et al. Estimating the causal tissues for complex traits and diseases. bioRxiv (2016).

14. Hu X. et al. Integrating Autoimmune Risk Loci with Gene-Expression Data Identifies Specific Pathogenic Immune Cell Subsets. Am. J. Hum. Genet. 89, 496–506 (2011).

15. Slowikowski, K., Hu, X. & Raychaudhuri, S. SNPsea: an algorithm to identify cell types, tissues and pathways affected by risk loci. Bioinformatics 30, 2496–2497 (2014).

16. Pers T. H. et al. Biological interpretation of genome-wide association studies using predicted gene functions. Nat. Commun. 6, 5890 (2015).

17. Gormley P. et al. Meta-analysis of 375,000 individuals identifies 38 susceptibility loci for migraine. Nat. Genet. 48, 856–866 (2016).

18. Fehrmann R. S. N. et al. Gene expression analysis identifies global gene dosage sensitivity in cancer. Nat. Genet. 47, 115–125 (2015).

19. Wood A. R. et al. Defining the role of common variation in the genomic and biological architecture of adult human height. Nat. Genet. 46, 1173–1186 (2014).

20. Locke A. E. et al. Genetic studies of body mass index yield new insights for obesity biology. Nature 518, 197–206 (2015).

21. Cahoy J. D. et al. A Transcriptome Database for Astrocytes, Neurons, and Oligodendrocytes: A New Resource for Understanding Brain Development and Function. J. Neurosci. 28, 264–278 (2008).

22. Akbarian S. et al. The PsychENCODE project. Nat. Neurosci. 18, 1707–1712 (2015).

23. Heng, T. S. P., Painter, M. W. & Immunological Genome Project Consortium. The Immunological Genome Project: networks of gene expression in immune cells. Nat. Immunol. 9, 1091–1094 (2008).

24. The 1000 Genomes Project Consortium. A global reference for human genetic variation. Nature 526, 68–74 (2015).

25. Sudlow C. et al. UK Biobank: An Open Access Resource for Identifying the Causes of a Wide Range of Complex Diseases of Middle and Old Age. PLOS Med 12, e1001779 (2015).

26. Okbay A. et al. Genome-wide association study identifies 74 loci associated with educational attainment. Nature 533, 539–542 (2016).

27. Okbay A. et al. Genetic variants associated with subjective well-being, depressive symptoms, and neuroticism identified through genome-wide analyses. Nat. Genet. 48, 624–633 (2016).

28. Teslovich T. M. et al. Biological, clinical and population relevance of 95 loci for blood lipids. Nature 466, 707–713 (2010).

29. Schunkert H. et al. Large-scale association analysis identifies 13 new susceptibility loci for coronary artery disease. Nat. Genet. 43, 333–338 (2011).

30. Manning A. K. et al. A genome-wide approach accounting for body mass index identifies genetic variants influencing fasting glycemic traits and insulin resistance. Nat. Genet. 44, 659–669 (2012).

31. Okada Y. et al. Genetics of rheumatoid arthritis contributes to biology and drug discovery. Nature 506, 376–381 (2013).

32. Jostins L. et al. Host-microbe interactions have shaped the genetic architecture of inflammatory bowel disease. Nature 491, 119–124 (2012).

33. Bradfield J. P. et al. A Genome-Wide Meta-Analysis of Six Type 1 Diabetes Cohorts Identifies Multiple Associated Loci. PLOS Genet 7, e1002293 (2011).

34. Dubois P. C. A. et al. Multiple common variants for celiac disease influencing immune gene expression. Nat. Genet. 42, 295–302 (2010).

35. Bentham J. et al. Genetic association analyses implicate aberrant regulation of innate and adaptive immunity genes in the pathogenesis of systemic lupus erythematosus. Nat. Genet. 47, 1457–1464 (2015).

36. Cordell H. J. et al. International genome-wide meta-analysis identifies new primary biliary cirrhosis risk loci and targetable pathogenic pathways. Nat. Commun. 6, 8019 (2015).

37. Anttila V. et al. Analysis of shared heritability in common disorders of the brain. bioRxiv 048991 (2016).

38. Lambert J.-C. et al. Meta-analysis of 74,046 individuals identifies 11 new susceptibility loci for Alzheimer’s disease. Nat. Genet. 45, 1452–1458 (2013).

39. Cross-Disorder Group of the Psychiatric Genomics Consortium. Genetic relationship between five psychiatric disorders estimated from genome-wide SNPs. Nat. Genet. 45, 984–994 (2013).

40. International League Against Epilepsy Consortium on Complex Epilepsies. Genetic determinants of common epilepsies: a meta-analysis of genome-wide association studies. Lancet Neurol. 13, 893–903 (2014).

41. Woo D. et al. Meta-analysis of genome-wide association studies identifies 1q22 as a susceptibility locus for intracerebral hemorrhage. Am. J. Hum. Genet. 94, 511–521 2014).

42. Traylor M. et al. Genetic risk factors for ischaemic stroke and its subtypes (the METASTROKE collaboration): a meta-analysis of genome-wide association studies. Lancet Neurol. 11, 951–962 (2012).

43. Patsopoulos N. A. et al. Genome-wide meta-analysis identifies novel multiple sclerosis susceptibility loci. Ann. Neurol. 70, 897–912 (2011).

44. Nalls M. A. et al. Large-scale meta-analysis of genome-wide association data identifies six new risk loci for Parkinson’s disease. Nat. Genet. 46, 989–993 (2014).

45. Ripke S. et al. Biological insights from 108 schizophrenia-associated genetic loci. Nature 511, 421–427 (2014).

46. Wain L. V. et al. Genome-wide association analyses for lung function and chronic obstructive pulmonary disease identify new loci and potential druggable targets. Nat. Genet. 49, 416–425 (2017).

47. Warren H. R. et al. Genome-wide association analysis identifies novel blood pressure loci and offers biological insights into cardiovascular risk. Nat. Genet. 49, 403415 (2017).

48. Lu Q. et al. Systematic tissue-specific functional annotation of the human genome highlights immune-related DNA elements for late-onset Alzheimer’s disease. PLOS Genet. 13, e1006933 (2017).

49. Tfelt-Hansen P. C. & Koehler P. J. One hundred years of migraine research: major clinical and scientific observations from 1910 to 2010. Headache 51, 752–778 (2011).

50. Backenroth D. et al. Tissue-specific functional effect prediction of genetic variation and applications to complex trait genetics. bioRxiv (2016).

51. Wilens, T. E., Biederman, J. & Spencer, T. J. Attention Deficit/Hyperactivity Disorder Across the Lifespan. Annu. Rev. Med. 53, 113–131 (2002).

52. Davis L. K. et al. Partitioning the Heritability of Tourette Syndrome and Obsessive Compulsive Disorder Reveals Differences in Genetic Architecture. PLOS Genet 9, e1003864 (2013).

53. Hanford, L. C., Nazarov, A., Hall, G. B. & Sassi, R. B. Cortical thickness in bipolar disorder: a systematic review. Bipolar Disord. 18, 4–18 (2016).

54. Callicott J. H. et al. Physiological Dysfunction of the Dorsolateral Prefrontal Cortex in Schizophrenia Revisited. Cereb. Cortex 10, 1078–1092 (2000).

55. Medic N. et al. Increased body mass index is associated with specific regional alterations in brain structure. Int. J. Obes. 40, 1177–1182 (2016).

56. Maleki N. et al. Migraine attacks the Basal Ganglia. Mol. Pain 7, 71 (2011).

57. Herculano-Houzel S. & Lent R. Isotropic Fractionator: A Simple, Rapid Method for the Quantification of Total Cell and Neuron Numbers in the Brain. J. Neurosci. 25, 2518–2521 (2005).

58. Sakai T. et al. Changes in density of calcium-binding-protein-immunoreactive GABAergic neurons in prefrontal cortex in schizophrenia and bipolar disorder. Neuropathology 28, 143–150 (2008).

59. Benes F. M. & Berretta S. GABAergic Interneurons: Implications for Understanding Schizophrenia and Bipolar Disorder. Neuropsychopharmacology 25, 1–27 (2001).

60. Gjoneska E. et al. Conserved epigenomic signals in mice and humans reveal immune basis of Alzheimer’s disease. Nature 518, 365–369 (2015).

61. Gagliano S. A. et al. Genomics implicates adaptive and innate immunity in Alzheimer’s and Parkinson’s. bioRxiv (2016). doi:10.1101/059519

62. Rege S. & Hodgkinson S. J. Immune dysregulation and autoimmunity in bipolar disorder: Synthesis of the evidence and its clinical application. Aust. N. Z. J. Psychiatry 47, 1136–1151 (2013).

63. Elamin, I., Edwards, M. J. & Martino, D. Immune dysfunction in Tourette syndrome. Behav. Neurol. 27, 23–32 (2013).

64. Jin, W., Millar, J. S., Broedl, U., Glick, J. M. & Rader, D. J. Inhibition of endothelial lipase causes increased HDL cholesterol levels in vivo. J. Clin. Invest. 111, 357–362 (2003).

65. Broedl U. C. et al. Endothelial lipase promotes the catabolism of ApoB-containing lipoproteins. Circ. Res. 94, 1554–1561 (2004).

66. Feingold K. R. & Grunfeld C. The role of HDL in innate immunity. J. Lipid Res. 52, 1–3 (2011).

67. Lo J. C. et al. Lymphotoxin beta receptor-dependent control of lipid homeostasis. Science 316, 285–288 (2007).

68. Harrison D. G. The Immune System in Hypertension. Trans. Am. Clin. Climatol. Assoc. 125, 130–140 (2014).

69. Hotamisligil G. S. Inflammation and metabolic disorders. Nature 444, 860–867 (2006).

70. Zlotnikov-Klionsky Y. et al. Perforin-Positive Dendritic Cells Exhibit an Immuno-regulatory Role in Metabolic Syndrome and Autoimmunity. Immunity 43, 776–787 (2015).

71. Dhirapong A. et al. B cell depletion therapy exacerbates murine primary biliary cirrhosis. Hepatol. Baltim. Md 53, 527–535 (2011).

72. Zhang J. et al. Ongoing activation of autoantigen-specific B cells in primary biliary cirrhosis. Hepatol. Baltim. Md 60, 1708–1716 (2014).

73. Raj T. et al. Polarization of the Effects of Autoimmune and Neurodegenerative Risk Alleles in Leukocytes. Science 344, 519–523 (2014).

74. Huang K. et al. A common haplotype lowers PU.1 expression in myeloid cells and delays onset of Alzheimer’s disease. Nat. Neurosci. 20, 1052–1061 (2017).

75. Lloyd C. M. & Hessel E. M. Functions of T cells in asthma: more than just TH2 cells. Nat. Rev. Immunol. 10, (2010).

76. Müller-Ladner, U., Pap, T., Gay, R. E., Neidhart, M. & Gay, S. Mechanisms of disease: the molecular and cellular basis of joint destruction in rheumatoid arthritis. Nat. Clin. Pract. Rheumatol. 1, 102–110 (2005).

77. Xavier R. J. & Podolsky D. K. Unravelling the pathogenesis of inflammatory bowel disease. Nature 448, 427–434 (2007).

78. Sospedra M. & Martin R. Immunology of Multiple Sclerosis. Annu. Rev. Immunol. 23, 683–747 (2005).

79. Barbosa, I. G., Machado-Vieira, R., Soares, J. C. & Teixeira, A. L. The immunology of bipolar disorder. Neuroimmunomodulation 21, 117–122 (2014).

80. Steiner J. et al. Acute schizophrenia is accompanied by reduced T cell and increased B cell immunity. Eur. Arch. Psychiatry Clin. Neurosci. 260, 509–518 (2010).

81. Sekar A. et al. Schizophrenia risk from complex variation of complement component 4. Nature 530, 177–183 (2016).

82. Corces M. R. et al. Lineage-specific and single-cell chromatin accessibility charts human hematopoiesis and leukemia evolution. Nat. Genet. 48, 1193–1203 (2016).

83. Ip H. et al. Stratified Linkage Disequilibrium Score Regression reveals enrichment of eQTL effects on complex traits is not tissue specific. bioRxiv 107482 (2017). doi:10.1101/107482

84. Mancuso N. et al. Integrating Gene Expression with Summary Association Statistics to Identify Genes Associated with 30 Complex Traits. Am. J. Hum. Genet. 100, 473–487 (2017).

85. Gusev A. et al. Integrative approaches for large-scale transcriptome-wide association studies. Nat. Genet. 48, 245–252 (2016).

86. Leeuw, C. A. de, Mooij, J. M., Heskes, T. & Posthuma, D. MAGMA: Generalized Gene-Set Analysis of GWAS Data. PLOS Comput Biol 11, e1004219 (2015).

87. Gazal S. et al. Linkage disequilibrium dependent architecture of human complex traits reveals action of negative selection. bioRxiv 082024 (2017). doi:10.1101/082024; Nature Genetics, in press

88. Boyle, E. A., Li, Y. I. & Pritchard, J. K. An Expanded View of Complex Traits: From Polygenic to Omnigenic. Cell 169, 1177–1186 (2017).

## References

89. Shi, H., Kichaev, G. & Pasaniuc, B. Contrasting the Genetic Architecture of 30 Complex Traits from Summary Association Data. Am. J. Hum. Genet. 99, 139–153 (2016).

90. Bulik-Sullivan B. K. et al. LD Score regression distinguishes confounding from polygenicity in genome-wide association studies. Nat. Genet. 47, 291–295 (2015).

91. Wagner, G. P., Kin, K. & Lynch, V. J. Measurement of mRNA abundance using RNA-seq data: RPKM measure is inconsistent among samples. Theory Biosci. Theor. Den Biowissenschaften 131, 281–285 (2012).

92. Law, C. W., Chen, Y., Shi, W. & Smyth, G. K. voom: precision weights unlock linear model analysis tools for RNA-seq read counts. Genome Biol. 15, R29 (2014).

93. Genovese G. et al. Increased burden of ultra-rare protein-altering variants among 4,877 individuals with schizophrenia. Nat. Neurosci. 19, 1433–1441 (2016).

94. Loh P.-R. et al. Efficient Bayesian mixed-model analysis increases association power in large cohorts. Nat. Genet. 47, 284–290 (2015).

95. Banda Y. et al. Characterizing Race/Ethnicity and Genetic Ancestry for 100,000 Subjects in the Genetic Epidemiology Research on Adult Health and Aging (GERA) Cohort. Genetics 200, 1285–1295 (2015).

96. Loh P.-R. et al. Contrasting genetic architectures of schizophrenia and other complex diseases using fast variance-components analysis. Nat. Genet. 47, 1385–1392 (2015).

97. Galinsky K. J. et al. Fast Principal-Component Analysis Reveals Convergent Evolution of ADH1B in Europe and East Asia. Am. J. Hum. Genet. 98, 456–472 (2016).

98. Chang C. C. et al. Second-generation PLINK: rising to the challenge of larger and richer datasets. GigaScience 4, 7 (2015).

